# The Role of Primate-Specific Genes and Network Dynamics in Cancer Evolution

**DOI:** 10.1101/2025.07.23.666306

**Authors:** Chuan Dong, Ya-Wei Su, Zhen Liao, Qian Xu, Hai-Xia Guo, Jianhai Chen, Wen Wei, Shengqian Xia, Zhenshun Cheng, Chengjun Zhang, Feng-Biao Guo

## Abstract

The atavistic evolutionary theory of cancer evolution highlights the role of ancient and unicell-originated genes, and comprehensive research demonstrated the evolutionary inference. Comparably, very few reports linked young genes with cancer development and maintenance. Here, we found that in cancer cell lines a higher proportion of core essential genes was present among primate-specific genes (PSG) than early metazoan-originated genes (Wilcoxon rank-sum test, *p-value* = 4.67e-12) and mammal-originated genes (Wilcoxon rank-sum test, *p-value* = 1.22e-16). Additionally, we found that the loss of many co-expression gene pairs in pan-cancer leads to their network becoming looser compared with normal tissue. However, PSGs particularly those being essential, exhibit a dynamic increase in co-expression, explaining their enhanced role in maintaining cancers. Furthermore, clustering of the gene co-expressions brings a cancer-exclusive module, which contains a large number of PSG connections. We also demonstrate the divergence between essential genes in healthy individuals and those in cancer cells. Our findings complement the atavistic theory in elucidating the evolutionary process of cancer.

## 1. Introduction

The evolution of complex multicellular organisms relies on highly regulated genetic networks and intercellular cooperation mechanisms, which ensure that the adaptability of individual cells is sacrificed in favor of achieving the optimal adaptability of the entire organism (Russo, et al. 2021). However, when affected by somatic mutations or other factors, these mechanisms may be disrupted, leading to cancer development. Several theoretical frameworks have been proposed regarding the origin of cancer. For example, the somatic mutation theory and the bad luck theory emphasize that the accumulation of somatic mutations leads to cell transformation, resulting in uncontrolled proliferation and, eventually, cancer development (Jassim, et al. 2023). The tissue organization field theory, on the other hand, proposes that malignant cancers arise from abnormalities at the tissue level. The ground state theory integrates elements from all three of these theories (Jassim, et al. 2023). Besides those, the atavistic evolution of cancer refers to cancer cells exhibiting characteristics or phenotypes similar to their unicellular ancestors during development and evolution (Boveri 2008; Chen, et al. 2015). During such a process, the evolution of cancer exhibits characteristics of convergent evolution in overall functional states (Chen and He 2016). The various theoretical frameworks on the origin of cancer illustrate the complexity of its developmental processes, which can be driven by endogenous or exogenous factors and nonpreventable or preventable processes (Cannataro, et al. 2022). Chen et al systematically carried out the experimental evolution of breast cancer using the xenografting method to characterize the complete evolutionary history of human breast cancer, which gave support to the theory of cancer atavistic evolution (Chen, et al. 2015). In addition, several other clues also illustrate the ancient origins of cancers. For example, cancer is widely present in many species, and cancer genes can be traced back to single-celled and early multicellular species (Domazet-Lošo and Tautz 2010). Regardless of the tissue origins of cancers, they consistently exhibit strong common characteristics in phenotypes, including unrestricted proliferation, abnormal response to signals, and uncontrolled cell growth. In essence, these shared traits suggest cancer cells change towards a unicellular-like state during their development (Hanahan and Weinberg 2011; Fouad and Aanei 2017), which can be as another evidence that cancer evolution is an atavistic evolution compared with species evolution. In a recent study, Trigos et al. showed that the distribution pattern of TAI (Transcriptome Age Index) and Gleason score (an index that can measure the malignancy of cancer) present a significant negative correlation, indicating that the enhanced expression of old genes facilitates the formation of cancer (Trigos, et al. 2017). Regarding atavistic reversions, Lineweaver et al. published a review paper that systematically summarizes the topic (Lineweaver, et al. 2021). The shared fundamental cellular functions required for the viability of all organisms are maintained by ancient genes (Miklos and Rubin 1996; Bergmiller, et al. 2012), which leads to the view that ancient genes typically associated with cancer development and some of them can become essential genes in pan-cancer (Bussey, et al. 2017).

Although atavistic evolution has received some supporting evidences, the atavistic evolution of cancer overlooks the impact of random genetic changes on cancer and other factors governing the development of cancer (Jassim, et al. 2023). Meanwhile, atavistic evolution also implies that newly formed genetic elements are less important compared to relatively ancient genes in cancer evolution, as these ancient genes are crucial for maintaining the activity of cancer cells. A recent study on pan-cancer co-expression networks revealed that many genes originating from multicellular organisms are activated in cancer and play crucial roles in tumor progression (Trigos, et al. 2024). Besides that, many *de novo* genes are involved in cancer because of the antagonistic co-evolution (McLysaght and Hurst 2016). Therefore, the contribution of multicellular-origin and *de novo* genes to the evolution of cancer should not be overlooked. Young/new genes refer to those genes that have recently emerged in the evolutionary process of a species. These genes retain the characteristics from the early stages of new gene birth, making them excellent materials for studying gene and phenotype evolution (Chen, et al. 2013). Within the atavistic evolution framework of cancer, ancient genes are suggested as potential targets for cancer therapy (Cisneros, et al. 2017; Russo, et al. 2021). Whereas young genes received little attention for the discovery of drug targets and hot spot genes in cancer evolution. Previously, the study of young genes mainly focused on their roles in adapting to the environment and promoting phenotypic innovation (Long, et al. 2013), while research on the functions of young genes in diseases has received relatively little attention. Recently, Ma et al. utilized data from The Cancer Genome Atlas (TCGA) project to demonstrate the cyclical nature of their involvement in cancer-related processes (Ma, et al. 2022). According to the theory of antagonistic pleiotropic (Williams 2001; Hughes, et al. 2002^)^, natural selection tends to select the beneficial effects of these genes, often overlooking their potential adverse consequences. Recent evidence demonstrated that young genes make contributions to disease phenotypes, highlighting their role in disease-associated phenotypic innovation (Chen, et al. 2013; Chen, et al. 2025). Considering the above reasons, these relatively young genes, such as primate-specific genes (PSGs), should receive more attention and investigations in the context of diseases especially cancers.

In this work, we proposed an index, pan-score, to measure a gene’s essentiality in pan-cancer, and depending on this we discovered that the PSGs can acquire cell-level essential functions in the process of cancer development. This was determined by assessing the proportion of cancer essential (CE) genes among the genes categorized by different gene origination times. Furthermore, our results suggest that the loss of co-expression pairs between genes in cancer leads to the pan-cancer network exhibiting a looser organization in comparison to the co-expression network observed in normal tissues (constructed by the gene expression from GTEx (Genotype-Tissue Expression) (Consortium 2020). Specifically, many gene pairs lost co-expression relationships during tumorigenesis, while the primate-specific genes acquired tighter co-expression relationships. In summary, these findings reveal that PSGs contribute to cancer development and have significant implications for cancer treatment. Such changes in gene co-expression can be summarized by the LinkLoss-Rec (co-expression link loss, and reconstruction) model, which successfully predicts the divergence in essential gene usage between human individuals and cancer cell lines.

## 2. Results

### 2.1 PSGs have a higher percentage of core CE genes compared to early metazoan genes and mammal-specific genes

We introduced and defined the pan-score, an index that could measure the essentiality of a gene in multiple cell lines and multiple cancer types (please refer to our method section for more details). The pan-score maintains high robustness when the number of sampled cell lines varies (Figure 1A). Particularly, when the number of randomly sampled cell lines increased to 40, the correlation *R* between the pan-scores calculated from all cell lines and those calculated from the sampled subset remains always over 0.95 (Figure 1A). Therefore, the pan-score could serve as a robust measure of the general essentiality of genes for the pan-cancer. Based on the analysis between the τ and the pan-score (Please refer to our method section for more details), our result illustrated that the CE (cancer essential) genes with higher pan-score tend to express in a wide range of cancer cell lines (R= –0.45 and *p-value* = 2.12e-274). However, this correlation decreased to –0.23 (R= –0.23 and *p-value* = 1.398e-10) for the correlation between the pan-score and the τ calculated by GTEx, implying that the expression pattern of pan-cancer is more correlated with the pan-score compared with the expression pattern of GTEx, and pan-score can reflect gene essentiality in pan-cancer instead of normal tissue. The pan-score for the essential genes of cancer cell lines is available from our Supplementary Table S1. Please note that in our work, ‘essential genes of cancer cell lines’ refers to the same set of genes as ‘cancer essential genes’.

**Figure 1.**
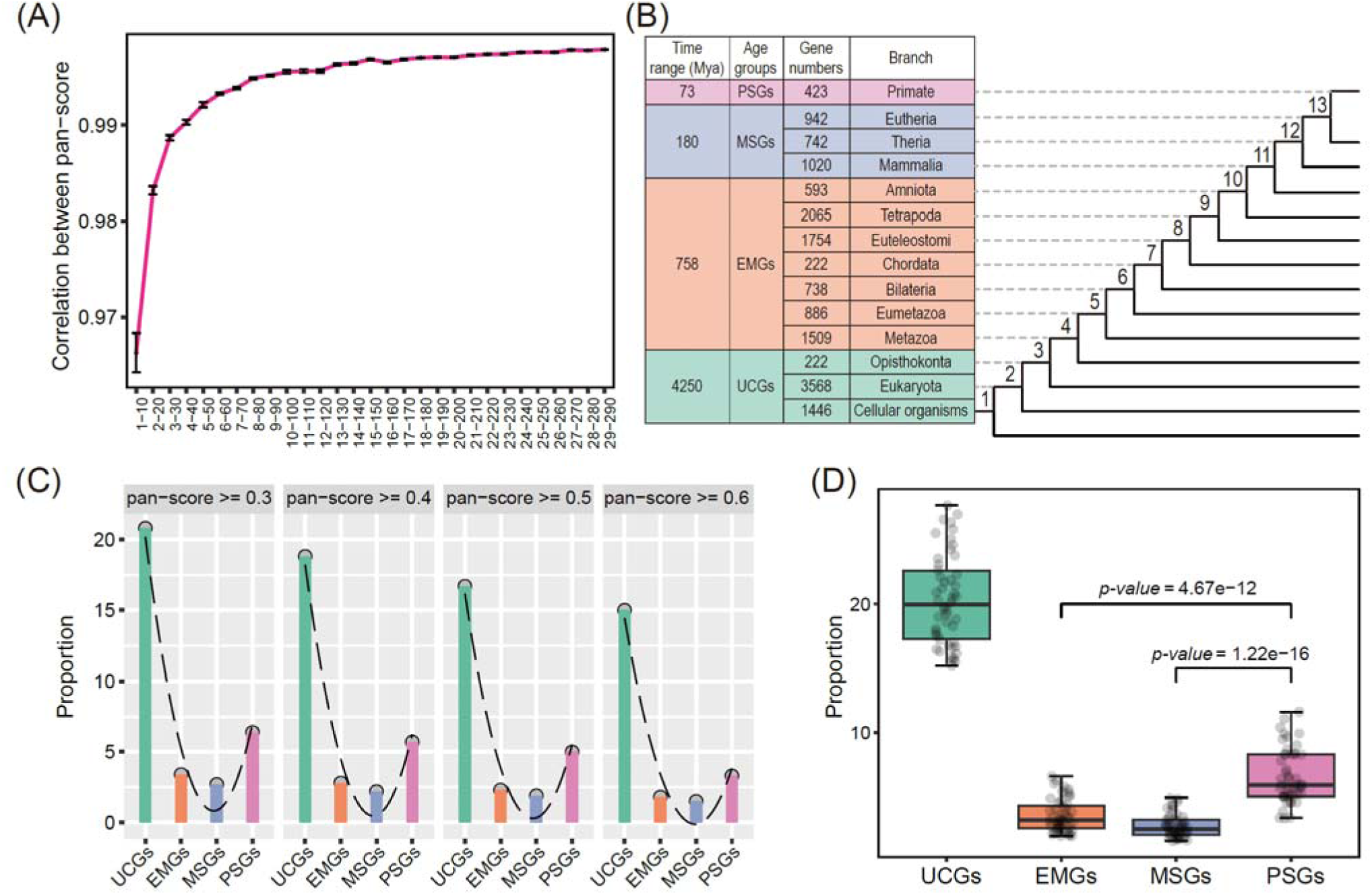
PSGs accumulate a higher percentage of core CE genes compared to early metazoan genes (EMGs) and mammal-specific genes (MSGs). A) The correlation analysis between pan-scores is based on all 323 cancer cell lines and those from the sampled cancer cell lines. As can be seen, the pan-score is a very robust measure of pan-cancer’s essentiality. The dot represents the mean correlation coefficient. In Figure A, the x-axis uses “Number1—Number2” to represent the data, where the first number indicates the round of simulation. In each round of simulation, Number2 means that Number2 samples were randomly selected: the number of cell lines sampled each time is indicated by Number2. For example, 29-290 means that in the 29^th^ round of simulation, 290 cell lines were randomly selected to re-calculate the pan-score, and this sampling was repeated 10 times in each round of simulation. The error bar is also indicated in Figure 1A. B) Summary information on gene ages, which was adapted from the results of Ma et al. (Ma, et al. 2022). There are four age groups that are highlighted by four different colors (each color corresponds to an age group), which can be further divided into 14 detailed age groups. The branch names are indicated adjacent to the branch on the evolutionary tree leading to primates. The number on the left of Figure 1B represents the time range for the four age groups, which is obtained from the TimeTree (the time scale of life) database (Kumar, et al. 2022). C) The percentage of core essential genes of cancer cell lines among the genes within the same age under different pan-score cutoffs. The pan-score used to define core essential genes is labeled at the top of each subfigure of Figure 1C. The x-axis represents four different age groups, while the y-axis denotes the proportion of core CE genes in genes with different origination ages. D) Core CE gene comparisons were conducted across multiple pan-score cutoffs (ranging from 0.1 to 0.6 with a step size of 0.01). Additionally, significance tests were performed between PSGs and MSGs, as well as between PSGs and EMGs.

Gene ages of humans could be classified into four broad-scale catalogs according to their origin time, named unicellular genes (UCGs), early metazoan genes (EMGs), mammal-specific genes (MSGs) and primate-specific genes (PSGs) (Figure 1B). The gene age data was adapted from the work of Ma et al.’s work (Ma, et al. 2022). We further divided the genes into two categories: core CE genes and non-core CE genes according to our proposed pan-scores. Genes with pan-scores above the threshold are referred to as core CE genes (Figure 1C), while those below the threshold are termed non-core CE genes. We changed the pan-score cutoff (pan-score ≥ 0.3, pan-score ≥ 0.4, pan-score ≥ 0.5 and pan-score ≥ 0.6) and calculated four ratios of core CE genes among the four gene catalogs divided according to their origination time. We found that the core CE genes in the primate-specific gene group present a higher percentage than the ones from EMGs and MSGs, despite a relatively lower percentage relative to the most ancient genes from unicellular ancestors (Figure 1C). After conducting a significance test (Wilcoxon rank-sum test) for the ratios with multi-pan-score cutoffs, we found that PSG’s ratios are significantly higher than those of EMGs and MSGs (*p-value* = 1.22e-16 between PSGs and MSGs; *p-value* = 4.69e-12 between PSGs and EMGs) (Figure 1D). Furthermore, if we adopt the ratio of core cancer essential gene number to the total number of CE genes, the difference becomes more significant (Supplementary Figure S1 in Supplementary Materials doc file). Previous extensive studies on ancient genes have revealed their critical roles in cancer development and species evolution (Miklos and Rubin 1996; Jordan, et al. 2002), which aligns with the predictions of the atavistic evolutionary theory of cancer. Based on this and the atavistic evolution theory of cancer, it is expected to observe gradually lower proportions of core CE genes among all genes as the birth of a new evolutionary branch. However, we found that the core CE genes in the primate-specific gene group present a higher percentage with statistically significant differences than the ones from EMGs and MSGs (Figure 1D). The core CE percentage does not change as a monotonic decrease when the gene’s origination time becomes recent, instead all four thresholds give parabolic patterns (Figure 1C). To further explore this, we checked if we could observe the same pattern using the small-scale age groups of 14 branches. Similarly, the core CE percentage in PSG is higher than those of the older age groups (branch 4 to branch 13) but lower than those of branches 1, 2 and branch 3 (which belong to UCGs) (Supplementary Figure S2 in Supplementary Materials doc file). Interestingly, there is a gradual decrease in the ratio of core CE genes as the gene’s origination time becomes recent (from cellular organisms to Chordata) (Supplementary Figure S2 in Supplementary Materials doc file). However, there is a sharp increase in the core CE gene ratio for the Tetrapoda-originated genes (Supplementary Figure S2 in Supplementary Materials doc file). In other words, the distribution of core CE genes throughout the evolutionary process from early metazoan species to primates does not depend on the age of gene origin, especially for the genes originating from Tetrapoda to primates (Supplementary Figure S2 in Supplementary Materials doc file). Furthermore, when we examined the distribution of essential genes (including core CE and non-core CE) of cancer cell lines across each branch, we did not observe the expected trend of a linear decrease in the proportion of essential genes with gene origin time become younger and younger, as would be anticipated in the theory of atavistic evolution (Supplementary Figure S3 in Supplementary Materials doc file). The atavistic evolution of cancer emphasizes the critical role of ancient genes (especially for unicellular genes and early metazoan genes; for example, most cancer-related genes originated in unicellular organisms and early metazoans (Domazet-Lošo and Tautz 2010)), overlooking the role of the recently originated genes in cancer. However, our results suggest an unexpected and neglected role of primate-specific genes in cancer function and maintenance.

Zinc-finger proteins have a wide variety of zinc-finger domains, which can be involved in the interactions with DNA and RNA and can be involved in several cellular processes such as cell migration, signal transduction and transcriptional regulation and so on (Cassandri, et al. 2017). Among the pan-cancer essential genes, 174 are associated with ZNF genes, accounting for approximately 2.3% (174/7470) of the total. For pan-cancer essential genes originating from different evolutionary branches, we expected approximately 59, 69, 16, and 3 ZNF-associated genes in the UCG, EMG, MSG, and PSG age groups, respectively. However, we observed only 23 and 6 ZNF-associated genes in the UC and EM groups. In contrast, the number of ZNF-associated genes in the MM and PSG branches was significantly higher than expected, with 76 and 39 observed ZNF-associated genes, respectively (Chi-square test, *P-value* = 4.86e-52 for the MM group and *P-value* = 6.97e-98 for the PSG group). This observation suggests that young ZNF-related genes may play a crucial role in maintaining essential cellular functions of cancer. The age-associated distribution analysis of pan-cancer essential genes revealed that during cancer evolution, PSGs (relatively young genes) can become increasingly essential, supporting the critical role of some relatively young genes. Recently, statistical data showed that 39 out of 83 de novo genes have been implicated in various cancers (Broeils, et al. 2023b; Xia, et al. 2025), which supports that the new birth gene could play an important role in cancer phenotype.

### 2.2 Through losing a great number of existing interaction pairs, the pan-caner co-express network becomes looser compared with the one in normal tissue

We downloaded co-expression data constructed by GTEx and TCGA from GeneFriends (Raina, et al. 2023) and employed the same approach used by GeneFriends to generate co-expression networks for protein-coding genes. This calculation relies on mutual rank to establish co-expression connections. Genes are considered co-expressed if they rank within the top 20 of each other’s co-expression profiles. According to this method, the pan-cancer co-expression network involves 70,823 interaction pairs and 16,980 genes, while GTEx involves 88,786 interaction pairs and 17,179 genes (the co-expression network in human individuals). Although the difference between the numbers of genes linked in two conditions is not significant, the difference in co-expression numbers is much higher among the gene number and cancer has fewer co-expression than the co-expression in normal tissue. Additionally, the mean degree in the pan-cancer network is 4, while the mean degree in GTEx increases to 5, illustrating that more genes are organized together in normal tissue (GTEx) compared to the pan-cancer. Therefore, we conclude that the pan-caner co-expression network becomes incompact compared with the GTEx co-expression network. Additionally, to further validate the above conclusion, we constructed a co-expression network using an alternative method, in which the network was constructed with Pearson’s correlation coefficient 0.5, 0.6, and 0.7 with p-value<0.05, respectively. We found that regardless of the thresholds that we employed, the number of edges in the GTEx co-expression network is always greater than that in the pan-cancer co-expression network (Supplementary Table S2). Similarly, the average degree is also higher in normal tissue compared to pan-cancer (Supplementary Table S2). Therefore, using the co-expression network constructed by Pearson’s correlation-based threshold, the network connection of protein-coding genes in normal tissues is denser than that in cancer tissues, consistent with the conclusion drawn from that constructed using the top mutual-rank-based method.

Since the pan-cancer network becomes loosened compared to the co-expression network of human individuals, it can be inferred that during the evolution of cancer cell lines, the acquisition of co-expression pairs should be less than the loss of co-expression interactions. In addition, the protein-coding genes can be divided into four categories based on their origin time according to the gene age list from Ma et al (Ma, et al. 2022) (Figure 1B) to further investigate this question. Accordingly, there are a total of 10 types of co-expression (Figure 2A). Among them, four types of co-expressions correspond to gene pairs within the same age group and six types of co-expressions indicate genes between different age groups. Upon analyzing the co-expression relationships, we discovered that the proportion of co-expression relationships established between PSGs within the pan-cancer network surpasses those that have been lost (Figure 2A), which indicates that there is a higher acquisition of co-expression relationships among PSGs (Figure 3A), even though co-expression relationships tend to be lost during the transformation from normal tissue to cancer. When counting the number of co-expression pairs, we also found that the number of co-expressions in PSGs within pan-cancer is significantly higher than that within GTEx (Figure 2B).

**Figure 2.**
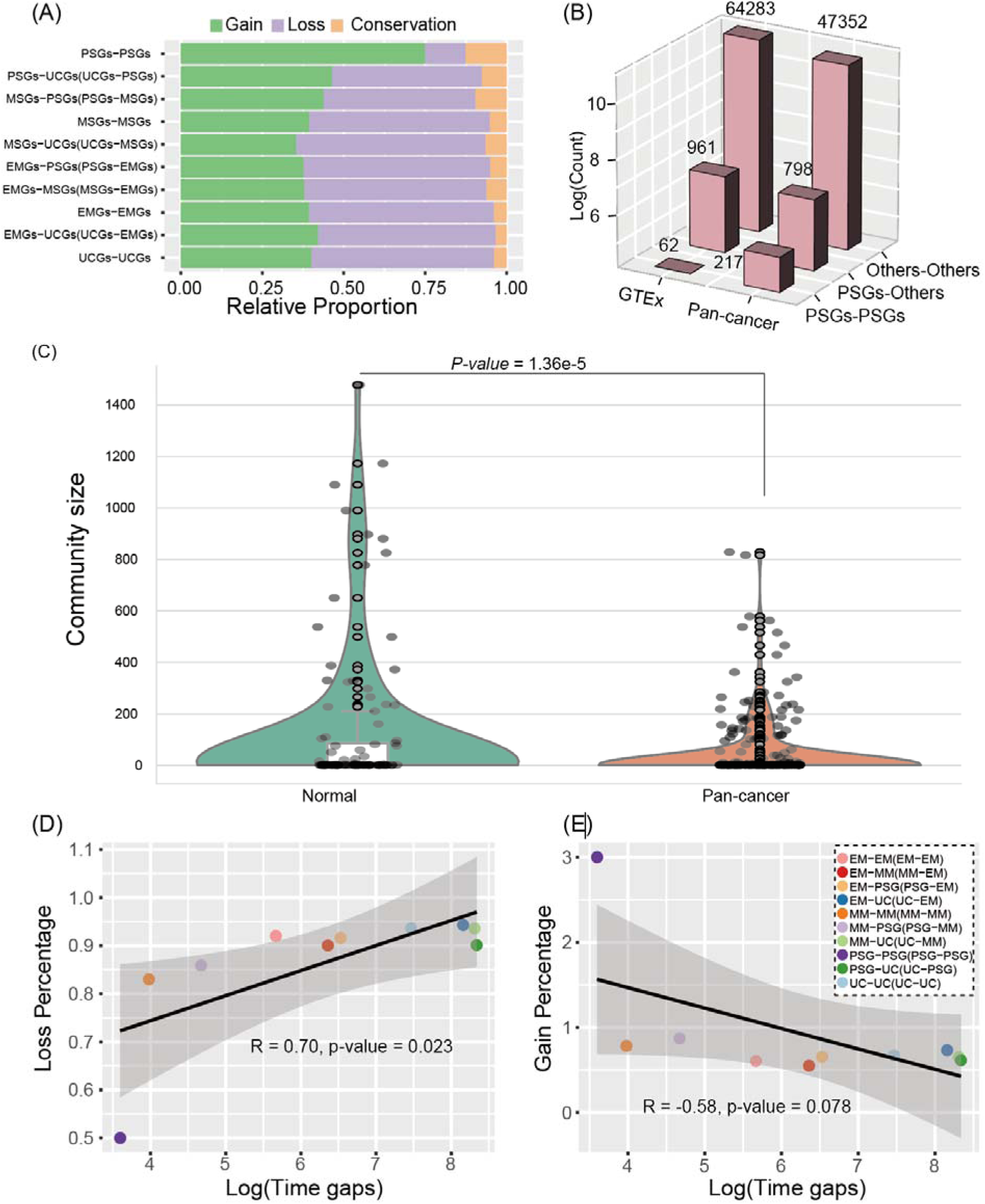
The comparisons between GTEx and pan-cancer co-expressions. A) The percentage of gain and loss in the co-expression of pan-cancer compared with the one in GTEx. B) The comparison of co-expression numbers between pan-cancer and normal tissue. The number of co-expressions from pan-cancer and GETx are indicated on the top of each bar. Please note that the z-axis uses a logarithmic scale. C). Size comparison of co-expression communities identified by the Leiden community detection algorithm. D) The loss of co-expressed relationship in pan-cancer (compared with co-expression in GTEx) varies with different intervals of origination time. The interval time was retrieved from TimeTree knowledge (Supplementary Figure S4 in Supplementary Materials doc file). Please note that we calculate the evolutionary time interval between a pair of genes within the same branch as half of the branch’s total time span. For example, the time interval for genes in PSG is 73/2 = 36.5 million years (MYs), while for genes in UCGs, it is 3492/2 = 1746 MYs (Supplementary Figure S4 in Supplementary Materials doc file). E) The gain of co-expression relationship in pan-cancer varies with the interval of origin between gene pairs. Figures 2D and Figures 2E share the same figure legend.

**Figure 3.**
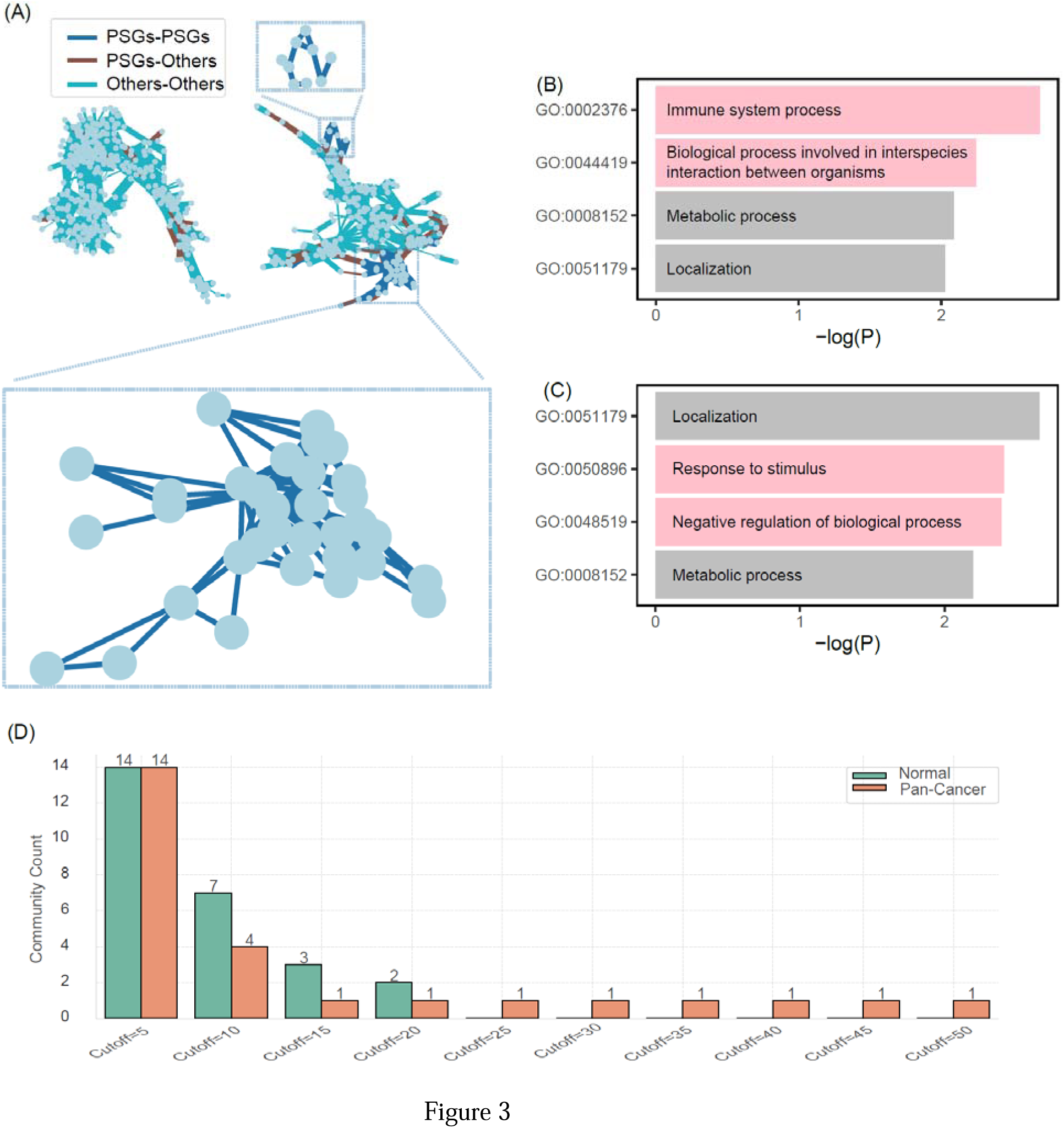
Co-expression clustering and functional enrichment analysis of CE and non-CE genes in PSGs, respectively. A) The comparisons are made for the sub-networks that contain the highest number of PSGs among all sub-clusters in pan-cancer and GTEx co-expression networks, respectively. We divided the co-expression links into three types, which contain the co-expression between PSGs and PSGs (represented by deep blue edges), the co-expression between PSGs and other genes (EMGs, MSGs and UCGs) (represented by brown edges) and the co-expression among other gene pairs (EMGs, MSGs and UCGs) (represented by cyan edges). The strength of co-expression is demonstrated by the thickness of the co-expression edges (reflected by their mutual rank). The light blue dashed rectangle in Figure 3A illustrates the enhanced co-expression module between PSGs. B) The enrichment analysis for the CE genes in PSG. C)The enrichment analysis for the non-CE genes in PSG. The red bars in (B) and (C) highlight the differences in enriched functions between primate-specific CE genes and primate-specific non-CE genes. D) Comparison of the number of communities containing more than a given number of PSGs.

The above result — that the normal co-expression network is denser than the overall pan-cancer interaction network — implies that the widespread loss of co-expression links in pan-cancer may lead to the formation of more communities compared with normal tissue. In other words, at the global level, the network communities (modules) in pan-cancer tend to become smaller. To validate this, we applied the Leiden community detection algorithm to identify network communities (co-expression modules) based on co-expression data (GenFriends) for normal tissue (GTEx) and pan-cancer (TCGA), respectively. Our results showed that 106 communities were detected in the co-expression network of normal tissue, whereas the number of communities in pan-cancer increased to 285, which is consistent with our inference. Furthermore, based on the detected network communities, we found that the average cluster size in normal tissue is larger than that in pan-cancer (Mann-Whitney U test, *P-value* = 1.36e-5) (Figure 2C). This pattern further supports our observation and is consistent with a recent study (Trigos, et al. 2024) showing that co-expression network modules constructed by WGCNA (Langfelder and Horvath 2008) across 33 pan-cancer types are smaller than those in normal tissue. From the global perspective of the entire co-expression network, our work together with that of Trigos et al. (Trigos, et al. 2024) demonstrates the sparsity of the pan-cancer network.

### 2.3 Originating-time-dependent loss and acquisition of gene co-expression relationships in the pan-cancer network, and the loss and acquisition of co-expression relationships exhibit opposite patterns

We wonder whether the change in co-expression (both loss and gain) is consistent across genes with different origination times. To explore this, we first defined co-expressed gene pairs that exist in the pan-cancer co-expression network but are absent in the normal tissue co-expression network as gained co-expressed gene pairs in pan-cancer. Similarly, we also defined lost co-expression gene pairs in the pan-cancer. Based on the pre-definition and our calculation, we examined the loss ratio of co-expression edges for gene pairs with different levels of evolutionary divergence. For a pair of genes within the same evolutionary branch, the time span of different branches varies. For example, PSG spans 73 million years (MYs) while the time of UCGs spans 3492 MYs. We represented the evolutionary time interval between a pair of genes within the same branch by using half of the branch’s total time span. For example, the time interval for a pair of genes in PSG is 73/2 = 36.5 MYs (Supplementary Figure S4 in Supplementary Materials doc file). According to this, we reveal that the trend of co-expression loss in pan-cancer exhibits an increasing trend as the divergent time increases (Figure 2D, R= 0.7, *P-value*=0.023). However, when we applied the same method to investigate co-expression gain, we observed a gradual decreasing trend, suggesting that co-expression relationships are initially formed with genes from closely related evolutionary branches (Figure 2E, R=-0.58, *P-value*=0.078) during the development of cancer. This reminds us of the evolution of new genes in protein interaction networks. For the pattern of new gene evolution in protein interaction networks, it has been reported that new genes initially interact at the periphery of the network after its birth and gradually integrate into the existing network over time (Zhang, et al. 2015). Furthermore, gene pairs with similar origination times tend to interact within protein-protein interaction networks (Capra, et al. 2010). Such patterns can be explained by differences in subcellular localization among proteins originating at different evolutionary times (Dong, et al. 2025). The co-expression gaining for genes in pan-cancer networks displays a similar pattern to that of protein interactions in the protein-protein interaction network after gene birth (Figure 2E); however, co-expression losing of gene pairs in pan-cancer networks exhibits an opposite evolutionary pattern compared with co-expression gaining (Figure 2D). The co-expression links between gene pairs of primate origin in the pan-cancer co-expression network are tighter, with the extensive establishment of co-expression relationships by gaining the highest co-expression ratio (Figure 2E). This reflects the important role of primate-specific genes in cancer, where the enhanced co-expression links among primate-specific genes might explain the high proportion of core CE genes in PSGs that we detected above (Figure 1C and Figure 1D).

### 2.4 The PSG co-expressed more tightly in the pan-cancer co-expression network compared to the GTEx co-expression network

We have indicated that the overall structure of the pan-cancer co-expression network has become sparser whereas the PSGs exhibit more co-expression links compared with the one in normal tissue. This provides us with motivation for further exploration of the functional cooperativity of PSGs in pan-cancer. Additionally, although the number of edges between PSGs has increased, is it possible for PSGs to form relatively strong co-expression modules?

We applied the Leiden algorithm (Traag, et al. 2019) to detect communities within the normal tissue and pan-cancer co-expression networks, respectively. In pan-cancer networks, 53 primate-specific genes (PSGs) were concentrated in a larger cluster, with PSGs comprising 23.6% of this cluster (Supplementary Table S3). In this cluster, there are 952 co-expression gene pairs. Among these, 246 co-expression relationships are involved by primate-specific genes. Out of the 246 pairs, 120 pairs are exclusively between PSGs, accounting for 48.8%. The remaining 126 pairs are involved by PSGs and other genes (including UCGs, EMGs, and MSGs). Specifically, among these 126 pairs, 120 pairs (48.8%) are between PSGs and MSGs, 3 pairs (0.1%) are between PSGs and EMGs, and 3 pairs (0.1%) are between PSGs and UCGs. This means that the age-closed gene pairs enhanced their co-expression relationships, which is consistent with our above observation (Figure 2E). However, when applying the same method and parameters to cluster the normal tissue co-expression networks, we did not find any cluster where PSGs exceeded the number of 25 genes. The most concentrated cluster contained only 23 primate-origin genes, making up merely 5.9% of the module, and the connectivity of these 23 PSGs is lower than that of the PSGs in pan-cancer. Hence, the module comparison analyses between pan-cancer and normal tissue illustrate that primate-specific genes become more tightly linked during cancer development. We further examined the co-expression strength of the cluster involving the 53 primate-specific genes (which can be represented by the thickness of the line in Figure 3A), and our results indicate that in the pan-cancer co-expression network, not only does the PSGs establish more co-expression edges, but the strength of the co-expression for PSGs also increases, forming a larger co-expression modules (Figure 3A) compared with the co-expression in normal tissue. This finding highlights the important role of PSGs in cancer. To validate that increased co-expression can enhance essentiality, we calculated the net co-expression gain, defined as the difference between co-expression gain and co-expression loss. Our results showed that among PSGs with net gains, 44.9% are essential genes. Whereas we merely expected 35% of genes in PSGs to be essential, the number of expected essential genes in PSG is significantly lower than the observed one (Chi-square test, *P-value* = 0.0118). The above result suggests that CE genes have higher co-expression links than non-CE genes. Based on the above analysis, we conducted that the CE genes in PSGs may have different functions compared to the non-CE genes in PSGs. When we defined a network community as PSG-related only if the number of PSGs within it exceeded a predefined threshold, we found that a relatively large number of PSGs tended to cluster within a single network community. This indicates that although some primate-specific genes become more dispersed in the pan-cancer network, another subset forms pan-cancer–specific network communities and interacts more frequently (Figure 3D). In other words, PSGs show a polarized pattern in pan-cancer: some become more scattered, while others become more clustered.

We further performed GO enrichment analysis using Metascape (https://metascape.org/). The results show that genes associated with the GO:0002376 function (immune system process) and the GO:0044419 function (biological processes involved in interspecies interactions) in normal tissue tend to become essential in cancer cell lines (Figure 3B and Figure 3C). For example, PF4 is an immune-related gene that can suppress the proliferation and cytokine release of activated human T cells (Fleischer, et al. 2002), which play an essential function in cancer. Meanwhile, PF4 is also involved in the process GO:0044419.

### 2.5 The divergence between IE (individual essential) genes and CE (cancer essential) genes

Given the alterations between the co-expression networks of normal tissue and pan-cancer, we became interested in exploring the similarities and differences between essential genes in healthy individuals and those in cancer cell lines. To investigate this, we compared the essential gene sets from human individuals (IE, from gnomAD) and pan-cancer cell lines (CE) across four age groups. The gnomAD deposited the aggregation of genetic variations in 125,748 exomes and 15,708 human individual genomes (Koch 2020), and defined the “loss of function tolerance” score based on the mutation frequency, which is named as LOEUF (loss-of-function observed/expected upper bound fraction). Genes with LOEUF values lower than 0.35 could be regarded as essential genes of human individuals (the cutoff is recommended by the official website), which typically experience purifying selection for those genes. We hypothesized that the loss and rewiring of co-expression links in pan-cancer would lead to distinct differences between these two essential gene sets. In this section, we tested this hypothesis using essential genes from gnomAD (IE) and essential genes of cancer cell lines (CE).

We examined the overlap of essential gene sets between cancer cell lines and human individuals. We observed a decreasing overlapping ratio between these two gene sets as the gene’s origination time became recent, indicating that a larger proportion of CE genes are non-essential in human individuals when restricted to younger genes (Figure 4A). This finding implied that there exists a separation (or usage divergence) for the two essential gene sets between essential genes in human individuals and those in cancer cell lines especially for the evolutionary young genes (such as PSGs). Considering that the unmatched essential gene number between human individuals and cancer cell lines might introduce estimation bias for this observation, therefore we set the pan-score cutoff and made the essential gene number approximately the same between CE and IE genes. In such datasets, we observed the same separation trend (Figure 4B), which again validates our hypothesis: the usage divergence between IE genes and CE genes. Additionally, we compared the essential gene ratios of each origin branch between human individual and pan-cancer. We found that the ratios of essential genes in human individuals and the ones in pan-cancer show a complementary pattern (Figure 4C). In detail, for human individuals, the proportion of essential genes shows a gradual decline in the lineage leading from metazoan-originated genes to primates (Figure 4C). However, this trend exhibits a turning point in pan-cancer essential genes, specifically starting from branch 7, where the proportion of CE genes begins to increase (Figure 4C). This further indicates that the evolutionary patterns of essential genes in pan-cancer differ from those in normal/human individuals. The distinct evolutionary pattern of the CE gene highlights the importance of primate-specific genes in the pan-cancer (Chen, et al. 2012). There are differences in the selection pressures on essential genes between normal individuals and cancer cells (Merlo, et al. 2006). In cancer, natural selection operates at the cellular level, driving the adaptive evolution of tumor cells (e.g., enhancing proliferation, invasion, and metastatic capabilities). In contrast, in human individuals, natural selection acts at the organismal level, aiming to improve survival and reproductive success. Based on these differences in selection pressures, we speculate that the distinct distribution patterns of essential genes in individuals versus pan-cancer essential genes may be related to effective population size (*N_e_*). Additionally, the cancer-specific essential genes also present a linear increase among the cancer essential genes (the number of pan-cancer specific essential genes/the number of CE genes) not only for the gene divided into 14 age groups but also for the genes divided into four age group (Supplementary Figure S5 in Supplementary Materials doc file). If there are some false-negative genes in the non-IE gene list according to the gnomAD annotation, it may affect the observed linearly increasing trend. In order to avoid this possible bias, we built a random forest model (please refer to section II in our Supplementary Materials doc file for more details). On the testing dataset, our results showed that the predicted positive examples had significantly lower LOEUF values (Wilcoxon rank-sum statistic, *p-value*=6.2e-35) compared to our predicted negatives (Figure 4D) (please refer to section II in Supplementary Materials doc file for how we construct the testing dataset). Thus, the random forest predictions exhibited agreement with the LOEUF in the gnomAD database (Karczewski, et al. 2020). We applied this model to cancer-specific essential genes (CE genes that do not overlap with IE genes) and predicted IE genes from them. The prediction results indicated a declining trend in the percentage of model-predicted IE among the cancer-specific essential genes as the genes’ origination time becomes recent (Figure 4E). Hence, the increasing trend of overlapping genes observed in Figure 4 A and B does not change. We can safely conclude that the divergence of two types of essential genes gradually becomes stronger with genes’ age becoming younger, and PSGs play an important role in cancer cells than in healthy individuals.

The segregation of IE and CE genes brings to mind the aforementioned results, the pan-score having a different correlation with normal and cancer expression patterns, respectively. This changes in gene expression patterns often reflect alterations or reductions in gene function (Kahlem, et al. 2004). However, the separation pattern of essential genes suggests that not only are there reductions in expression and functional degradation, but also the essentiality of many genes’ functions changes when cancer cells emerge in normal human individuals. Additionally, the separation may also imply the metabolic reprogramming of cancer cell lines compared to normal tissues. Because we observed the increasing separation trend for the essential genes as the gene’s origination time became recent, we inferred that the PSGs may also be involved in metabolic reprogramming process.

Based on the separated utilization of essential genes (Bartha, et al. 2018), especially in PSGs between human individuals and cancer cell lines, we proposed that PSGs can also be utilized as targets for cancer therapy because most of these essential genes in cancer are non-essential in human individuals even when we apply a random forest model to check the conclusion. We checked the distribution of 148 cancer-specific essential PSGs in terms of LOEUF, pan-score and decision value from our RF-based model. We found that three genes are predicted as essential for human individuals (Supplementary Table S4). We proposed that these 145 rest genes with larger pan-score and larger LOEUF (the gene in the upper right corner of Figure 4F) can be utilized as targets in cancer treatment (Figure 4F). For example, the primate-specific gene ZNF730 (ENSG00000183850) emerges as a hub node with a degree of 8, indicating its significance as an essential gene across various cancers. However, according to gnomAD, this gene is classified as non-essential in individual humans. Furthermore, a random forest-based model validates its non-essential status in human individuals, assigning it a decision value of 0.03. Besides that, many Zinc finger genes become essential genes in pan-cancer (Supplementary Table S1), and the Zinc finger genes originated from primates also involved in the pan-cancer specific co-expression network. We therefore think that the primate-originated Zinc finger genes can be as preferred targets for cancer treatment. A recent work illustrates that cancer cells can employ zinc finger proteins to suppress TE-originating surveillance mechanisms, which may facilitate clonal expansion, diversification, and immune evasion (Martins, et al. 2024). This conclusion from the previous work (Martins, et al. 2024) highlights the crucial role of zinc finger genes in cancer maintenance, meanwhile our work in this section highlights the importance of primate-specific Zinc Finger genes.

## 3. Discussion

The successful treatment of cancer requires an understanding of the biological origin mechanisms within diseased tissues. Theories on the origins of cancer have provided valuable insights for its treatment. For instance, the atavistic evolutionary theory of cancer emphasizes the significant role of early-origin genes in cancer progression, suggesting that ancient essential genes could serve as potential therapeutic targets (Davies and Lineweaver 2011) and potential prognostic biomarkers (Kang, et al. 2025). However, this theory downplays the contributions of young/new genes, such as primate-specific genes in cancer. With the increasing availability of gene essentiality data from multiple cancer cell lines (Tsherniak, et al. 2017), it has become possible to systematically assess the essentiality of each gene at a genome-wide scale, Meanwhile, large-scale pan-cancer gene inactivation data provides a significant opportunity for the extensive testing of clinically viable drug targets (Blomen, et al. 2015; Hart, et al. 2015; Wang, et al. 2015; Chang, et al. 2021). A recent research has shown that genes of multicellular origin and genes of unicellular origin exhibit significant dynamic changes in co-expression networks during cancer evolution (compared to the co-expression patterns within unicellular-origin genes or within multicellular-origin genes) (Trigos, et al. 2024), indicating that the rewiring of co-expression between multicellular and unicellular genes play an important role in cancer. Such binary classification of genes into multicellular and single-cell origin categories, to some degree, overlooks the role of evolutionarily young genes in cancer development. Therefore, analyzing the proportion of cancer-essential genes across genes with different evolutionary origins could provide deeper insights into whether newly emerged genes also play indispensable and non-negligible roles in cancer. In this work, we utilized the most recently updated gene age data to divide genes into four detailed age groups, which can be further divided into 14 more detailed age groups (Ma, et al. 2022). Previous work showed that the integration of young genes into gene-gene interactions is a stepwise process (Zhang, et al. 2015) in which newly evolved genes preferentially interact with genes with similar age (Capra, et al. 2010). Our results illustrated that such newly formed links between young and relatively old genes are not stable, leading that they are easy to be lost under an adverse environment (Figure 2D). Furthermore, our result demonstrated that the loss of co-expression is a selective process, which implied that the loss of interaction pairs is a reverse process of young gene integration (Figure 2D and Figure 2E). In other words, during cancer evolution, genes tend to lose co-expression connections with those of distinct evolutionary origins and gain co-expression with genes of similar evolutionary time. Considering that young/new genes usually have stronger dynamics, plasticity and faster evolutionary speed (Cai and Petrov 2010; Vishnoi, et al. 2010; Tautz and Domazet-Lošo 2011), they should be more sensitive to environmental changes and able to adapt to them more quickly.

Cancers occur widely across the tree of life (with different species exhibiting varying degrees of susceptibility) (Vincze, et al. 2022; Compton, et al. 2025). This widespread presence can be well explained by the atavistic evolution of cancer. However, it cannot explain the high proportion of core CE genes appearing in PSG, which was found in our present work (Figure 1C and Figure 1D). According to the atavistic evolution, the enhancement of ancient gene functions and the loss of new gene functions is our anticipated phenomenon. Therefore, the observation that recently emerged PSGs have a higher proportion of essential genes than MSGs is, to some extent, inconsistent with the expectation of atavistic evolution. Based on our results, we summarize the cancer evolutionary process as link loss and reconstruction (LinkLoss-Rec) within the gene co-expression network (Figure 5), which may help explain this apparent contradiction. It emphasizes the loss and reconstruction of co-expression relationships during the evolution of cancer. This loss creates opportunities for the establishment of new co-expression regulatory networks among genes that previously had no co-expression relationships. Many co-expression links in normal tissue are lost during cancer development, particularly for gene pairs with distinct evolutionary origins (Figure 2D). As a result, some of the link-lost genes may re-establish their co-expressed relationships, especially among gene pairs with similar origins (Figure 2E). During this process, some PSGs enhanced their co-expression and established collaborative modules, which may contribute to their potential to become essential genes. Studies have shown that aggressive perturbations of network hub genes by somatic mutations can drive cells into new phenotypic states (Cheng, et al. 2014), such as tumorigenesis, highlighting the importance of network rewiring in maintaining additional phenotypic states. Evidence in the literature supports this deduction. For example, some young genes birthed by *de novo* can promote cancer development if such genes are aberrantly expressed in somatic cells (such expression pattern disrupt their original co-expression network) (Broeils, et al. 2023a). The created new co-expression module among PSGs might maintain the additional phenotype of cancer (IE genes and CE genes show the greatest divergence in PSGs). Disrupting the newly formed co-expression networks involving PSGs could aid cancer treatment, suggesting that this model may also offer valuable insights for therapeutic approaches. Furthermore, according to our result and LinkLoss-Rec, we proposed that cancer evolution is not a simple process of atavistic evolution, but the evolution of cancers is accompanied by dynamic changes in the co-expression network, especially for the interactions between PSGs and PSGs themselves. We are aware that it is certainly not the whole story of cancer, and many cancer-related processes, such as angiogenesis, immune evasion and tissue infiltration, may rely on evolutionary innovations rather than simple degeneration (Benton, et al. 2021). Separation of such innovative processes from degenerative processes will be critical for understanding the development of cancer disease and their treatment.

**Figure 4.**
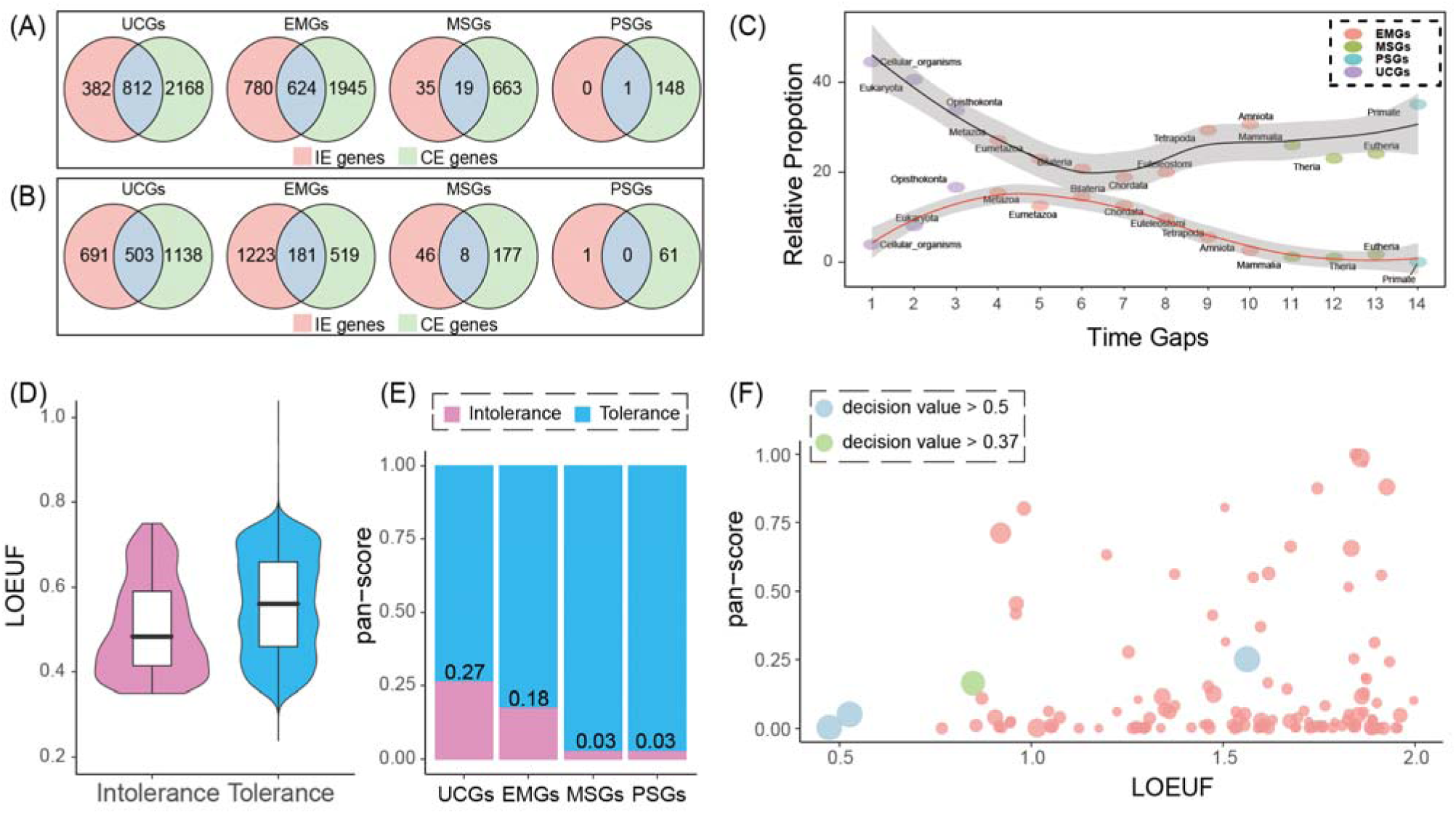
The relationship between IE and CE genes. A) The comparison of CE genes and IE genes in different age groups. It shows the decreasing ratio and decreasing number of overlapping genes between two essential gene categories (IE genes and CE genes). B) To avoid the statistical bias from the difference between the numbers of two gene catalogs (CE genes and IE genes), we change the pan-score cutoff and make the number of CE genes equal to the number of IE genes, then perform the same analysis as shown in (A). C) Comparison of essential gene ratios at different origin times between human individuals and pan-cancer, and the two curves exhibit a different pattern. D) The comparison of LOEUF scores between IE and non-IE genes predicted by our random forest model on the testing dataset. The Wilcoxon rank-sum statistic demonstrated a significantly lower LOEUF for the predicted positives compared with the predicted negatives (*P-value*=6.2e-35). E) The percentage of predicted IE genes in four categories (the ratio equals the number of predicted IEs within the unique CEs to the total number of unique CEs. The percentage is marked at the top of the red bar. F) The distribution for the 148 cancer-specific essential genes (Supplementary Table S4). It displays the results of random forest prediction for the 148 CE genes, where the circle size corresponds to the decision value in the random forest model.

**Figure 5.**
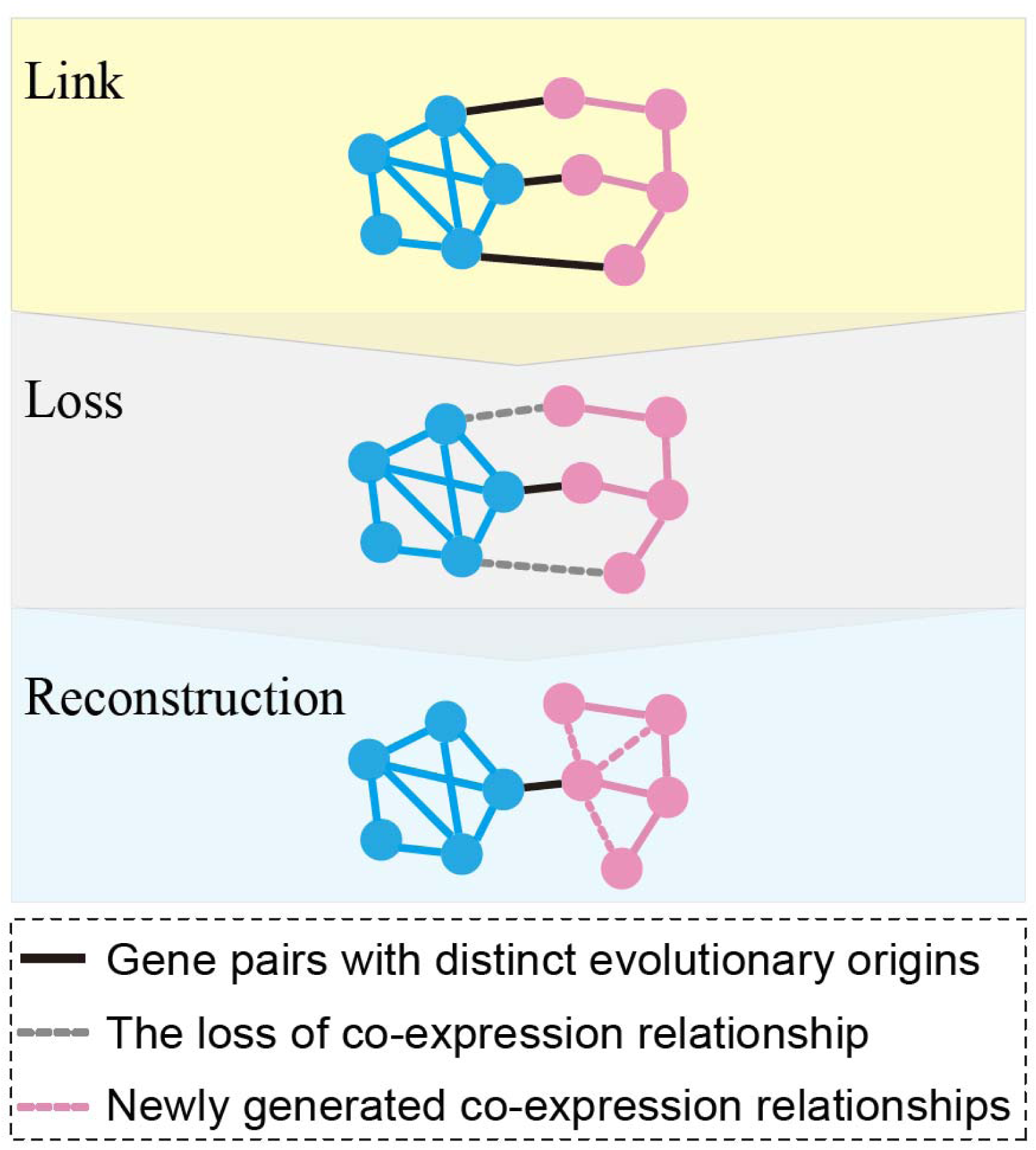
The link loss and reconstruction (LinkLoss-Rec) model. Dashed gray line: the interaction formed by genes with distinct evolutionary origination. Dashed red line: the newly formed co-expression links, which are usually connected by the gene pairs with closed evolutionary origination. The blue solid line: The unchanged co-expression relationship between the co-expression network of normal tissue and the co-expression network of pan-cancer.

Our findings reveal a shift in essential gene usage between healthy individuals and pan-cancer samples. During cancer progression, many genes lose their original co-expression partners and form new connections, causing previously peripheral genes to emerge as network hubs in the pan-cancer co-expression network. This rewiring alters which genes are essential, creating a clear separation between the gene sets of normal and cancer tissue. PrimateLspecific genes (PSGs), which undergo dynamic rewiring during this process, display pronounced shifts in essentiality. These observations underscore the pivotal role of network remodeling in redefining gene essentiality in pan-cancer. Based on the essential gene segregation in PSGs (Figure 4A and Figure 4B), we speculate that the young genes can serve as preferred drug/vaccine targets for cancer treatment. For example, LGALS9C is a primate-specific gene and exhibits tissue-specific expression. It has a pan-score value of 0.315 and demonstrates essential functions in several cancer cell lines. It is worth noting that this gene has already started to play a role in cancer treatment (Lv, et al. 2022). In this work, we identified approximately 140 cancer therapeutic target genes that are primate-specific (Supplementary Table S4). This was achieved through screening using random forest models, the gnomAD database, and essential gene data from cancer cell lines. Those PSGs can form pan-cancer-specific co-expression sub-networks. Therefore, the hub genes in this pan-cancer-specific sub-network can also be considered as candidate targets for cancer treatment. We also check the newly formed PSG-PSG co-expression pairs. We found that many genes can encode ZNF proteins, which can regulate cell migration (Cosby, et al. 2021), implying their role in cell movement. Some zinc finger proteins have essential functions in cancer (Ying, et al. 2022; De Franco, et al. 2023), which is consistent with our inference. In addition, such genes show differences in co-expression intensity between cancer and normal tissues.

## 4. Materials and methods

### 4.1 The dataset preparation for gene age, essential genes, and co-expression relationship

We downloaded the gene age data from the supplementary table of Ma et al’s work, where the gene ages were classified into 14 fine-scale phylogenetic stages and four broad-scale evolutionary epochs (Ma, et al. 2022). The Figure 1B provides a detailed summary of the genes’ age (Ma, et al. 2022) and their distribution in different origination branches.

The genes’ mutation tolerance data were downloaded from the gnomAD database, and genes with a LOEUF score lower than 0.35 were considered as essential genes of human individuals (IE genes), which is set according to the officially recommended cutoff. We obtained essential genes of cancer cell lines associated with 32 cancer types, including more than 300 cell lines, and the originated data can be retrieved from the supplementary data provided by Behan et al (Behan, et al. 2019).

The co-expression rank of normal tissue and pan-cancer was retrieved from GeneFriends knowledgebase (Raina, et al. 2023). The co-expression networks of pan-cancer and normal tissues were further constructed based on mutual rank according to the description at https://coxpresdb.jp/static/help/mr.shtml. We considered the gene pairs to be co-expressed when the mutual rank is within the top 20.

### 4.2 Calculating the pan-score for estimating gene essentiality in pan-cancer

CRISPR-Cas technology generated high-through essential gene data associated with cancer cell lines (Behan, et al. 2019), and information on essential genes has been curated by several online resources such as OGEE, DEG (Gurumayum, et al. 2021; Luo, et al. 2021) and DepMap (Tsherniak, et al. 2017). Considering that the heterogeneity between cell lines of the same cancer type is smaller than the heterogeneity between cell lines of different cancer types, a simple proportional measurement of the essentiality of cancer essential genes will introduce bias estimation into the calculation of gene’s essentiality in pan-cancer. Therefore, it is unreasonable to measure the essentiality of a gene solely based on the proportion of cell lines in which it is essential. To overcome this, we introduced and defined the pan-score, an index that can be used to measure the general essentiality of a gene in multiple cell lines and multiple cancer types, which can be calculated by the formula (1), where n represents the number of cell lines in the *j^th^* cancer type where gene *i* is an essential gene, and N represents the total number of cancer cell lines in the *j^th^* cancer type. The *N* represents the total number of cancer types. In our study, we have a total of 32 cancer types; hence, *N* is equal to 32. We have a total of 7010 cancer essential genes; therefore, *j* ranges from 1 to 7010 (representing the 7010 CE genes). Accordingly, the greater the pan-score value, the greater the corresponding pan-cancer essentiality, and the smaller the pan-score value, the smaller the corresponding essentiality (Supplementary Table S1). Additionally, we can define these genes as essential genes of pan-cancer if the pan-scores of these genes are larger than the users’ pre-given cutoff.

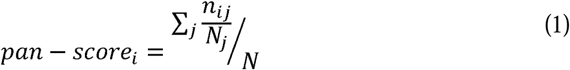

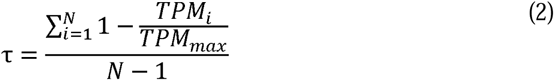

The calculation of the pan-score relies solely on our downloaded essential gene lists in more than 300 cancer cell lines. To ensure the robustness of the pan-score index across different datasets, we performed a series of Monte Carlo simulations by randomly sampling cancer cell lines from the more than 300 cancer cell lines to construct various datasets. Then, we recalculated the pan-score using these newly generated datasets. Based on all datasets, we can compute the initial pan-score, while based on each randomly sampled list, we can compute the pan-score again based on the randomly sampling cancer cell lines. The Pearson correlation coefficient can then be calculated between the pan-score of the overall samples and the pan-score of the randomly drawn samples. If a strong correlation exists between these two pan-scores calculated from different cancer cell samples, it could indicate that the pan-score can serve as a reliable index for measuring gene essentiality in pan-cancer research. For different numbers of cell lines, we conducted 10 random samplings and compared the Pearson correlation coefficients of two pan-scores in each round simulation (pan-score calculated based on the original sample versus pan-score calculated based on the random samples).

To assess the expression specificity in these cell lines, we utilized the τ index proposed by Yanai et al. (Yanai, et al. 2005). In formula (2), *N* represents the number of tissues, *TPM_i_*represents the gene expression level in the *i^th^* cancer cell line. The τ index ranges from 0 to 1, reflecting the expression degree of tissue specificity for a gene. Specifically, a high τ value indicates a narrow expression across tissues, whereas a low τ value indicates a broader expression across tissues.

### 4.3 The estimation for the gain and loss of co-expression links

We check the same pair of genes in both the GTEx co-expression network and the pan-cancer co-expression network, respectively. If a gene pair is present in the GTEx co-expression network but not in the pan-cancer co-expression network, we consider that this co-expression relationship has been lost during cancer development. Conversely, we can also identify newly formed (gained) co-expression relationships during cancer development, which are those co-expressed gene pairs present in the pan-cancer network but absent in the GTEx network. To further assess the extent of co-expression loss and gain during the transition from normal tissue to cancer, we use the percentage of lost or gain edge connections relative to the total number of GTEx edge connections as a metric.

### 4.4 Screening the false negatives from human individual non-essential genes by the random forest model

The genetic variations for some genes may lead to phenotypic effects occurring later in life, especially for the variation during post-reproductive years. Therefore, such genes should have a relatively lower LOEUF score (Loss-of-function observed/expected upper bound fraction) (Gudmundsson, et al. 2022), however, gnomAD might confer them a relatively higher LOEUF score. Based on this consideration, some false negatives exist in the non-human individual essential genes (for example, false negatives may exist among cancer-specific essential genes). To make an accurate analysis, we utilized the random forest model (performed in scikit-learn python package (Pedregosa, et al. 2011)) to identify such genes. To select genes for training, we used genes with a LOEUF cutoff lower than 0.35 (this cutoff is recommended by the gnomAD website version 2) as positive training data, which we considered such genes as essential genes of human individuals and genes with a LOEUF cutoff higher than 0.75 as negative training data (based on the data in gnomAD version 2). These genes are considered as non-essential genes of human individuals. The remaining genes with LOEUF ranging from 0.35 to 0.75 in gnomAD were used as a testing dataset (based on the data in gnomAD version 2). In summary, we have a training dataset including genes with LOEUF lower than 0.35 and genes with LOEUF larger than 0.75. Besides this, we also have a testing dataset with LOEUF between 0.35 and 0.75. The gene expression data from GTEx was used as the major feature to train the random forest model. Our proposed sequence features were also incorporated into account during our training and predicting process (Guo, et al. 2017; Dong, et al. 2020) (Section IV in our Supplementary Materials). Herein, we have two types of essential gene sets. One is derived from the cancer cell lines, referred to as essential genes of cancer cell lines (CE genes), and the other from the human individuals, known as human individual essential (IE) genes.

## Author contributions

Feng-Biao Guo designed this work. Feng-Biao Guo, Chengjun Zhang and Chuan Dong coordinated this work. Chuan Dong and Ya-Wei Su and Zhen Liao performed the computational analysis. Ya-Wei Su also double-check the main results. Chuan Dong and Zhen Liao drafted the manuscript, Jianhai Chen, Wen Wei, Zhen Liao, Hai-Xia Guo, Qian Xu, Zhenshun Cheng, and Shenqian Xia involved discussions and contributed to writing. Feng-Biao Guo, Chengjun Zhang, Jianhai Chen and Hai-Xia Guo further revised the manuscript.

## Supplementary Code

All the scripts used for generating the figures is available at figshare https://figshare.com/articles/figure/_b_Scripts_for_generating_data_and_figures_b_/28544765?file=52821824.

## Acknowledge

The Genotype-Tissue Expression (GTEx) Project was supported by the Common Fund of the Office of the Director of the National Institutes of Health and by NCI, NHGRI, NHLBI, NIDA, NIMH, and NINDS. The data used for the analyses described in this manuscript were obtained from the expression file named GTEx_Analysis_2017-06-05_v8_RNASeQCv1.1.9_gene_median_tpm.gct, which is downloaded on May 2023. We are grateful to Priyanka and Pedro for their assistance in resolving our questions about GeneFriends. We appreciate the valuable discussions with members of the Guo lab, and Dr. Zhigang Han. We gratefully acknowledge Manyuan Long for his constructive suggestions and insightful discussions.

